# Fibronectin 1 is required for suture patency and dysregulated across craniosynostosis models in the mouse

**DOI:** 10.64898/2026.03.04.709647

**Authors:** Xiaotian Feng, Megan Gregory, Alex Flores, Morgan Horowitz, Miranda Sun, D’Juan Farmer, Matthew P. Harris, Greg Holmes, Radhika P. Atit

## Abstract

The mammalian skull roof is comprised of calvarial bones, connected through fibrous sutures, that protect the brain and allow for growth. Premature suture closure, or craniosynostosis (CS), impedes expansion, impacting 1 in every 2500 newborns. Despite its genetic heterogeneity, with nearly 80 associated genes, CS manifests as a common phenotype, hinting at a convergent etiological mechanism. Recently, we described how graded expression of extracellular matrix protein Fibronectin (FN1) is required for apical expansion of calvaria and coronal suture patency. Dysregulated FN1 expression has been identified in two human CS syndromes, suggesting its potential as a convergent mechanism of CS. Here, we further investigate the cellular basis for the CS phenotype in the *Fn1* mutant mouse. Graded expression of FN1 baso-apically in the cranial mesenchyme was variably dysregulated across mouse models of syndromic and non-syndromic CS and accompanied by diminished apical expansion of frontal bone primordia. In parallel, at later developmental stages we find ectopic osteogenic induction of *Six2*^+^ patent suture mesenchyme in the *Fn1* mutant. These findings pinpoint FN1 as a crucial regulator of suture patency by modulating calvarial growth and driving cell identity, and differentiation, thus providing a potential target for matrix-mediated treatments.

## Introduction

The mammalian calvaria consists of five bones that form the roof of the skull and encase the brain. They are derived from both neural crest and cephalic mesoderm (Jiang et al., 2002; Yoshida et al., 2008). As the calvarial bones develop, fibrous joints known as sutures form between them, in which the growing edges of the bones, or osteogenic fronts, are separated by suture mesenchyme (Ishii et al., 2015; Jiang et al., 2002; Morriss-Kay and Wilkie, 2005). These sutures allow for the growth and expansion of the skull to accommodate the growing brain during prenatal and postnatal stages (White et al., 2021). In embryonic development, the growth of calvarial frontal and parietal bones initiates at foci above the eye in the supraorbital arch mesenchyme (Deckelbaum et al., 2012; Tran et al., 2010; Yoshida et al., 2008). Maintenance of a non-ossified suture mesenchyme is critical for normal calvarial growth (White et al., 2021). Studies over the years have identified many genes whose dysregulation perturbs calvarial growth, resulting in congenital calvarial defects such as craniosynostosis and various calvarial dysplasia.

Craniosynostosis (CS) is a congenital condition characterized by the premature fusion of one or more cranial sutures, leading to abnormal skull shape and potential developmental issues. There are a variety of common non-syndromic and syndromic human diseases with CS, which in total occurs in 1 in 2,500 newborns (Wilkie, 1997; Wilkie et al., 2017). The most common form of CS involves the fusion of a single suture, typically the sagittal suture, but syndromic CS usually involves the fusion of multiple sutures (Cunningham et al., 2007; Johnson and Wilkie, 2011). CS is not only a structural anomaly of the suture, but also involves complex biological processes and adversely can impact calvarial growth (Flaherty et al., 2016; Morriss-Kay and Wilkie, 2005; Teng et al., 2018; Twigg and Wilkie, 2015)

The activity of various growth factors, transcription factors, and signaling pathways are implicated in CS. Mutations in genes such as *FGFR1-3, TWIST1*, and *EFNB1* have been associated with syndromic CS which particularly involve the coronal suture(Passos-Bueno et al., 2008; Twigg and Wilkie, 2015; Wilkie et al., 2010). Transcriptomic analysis of single suture CS from a panel of well-characterized clinical samples identified extracellular matrix (ECM)-mediated focal adhesions to be differentially regulated in the different sutures of CS patients compared to control sutures(Stamper et al., 2011). There are over one hundred well established syndromic forms of CS with known genetic mutations (Aleck, 2004; Ishii et al., 2015; Kutkowska-Kaźmierczak et al., 2018; Timberlake et al., 2019; Wilkie et al., 2017). Some of the genes associated with non-syndromic CS are known and overlap with genes involved in syndromic CS, yet the etiology of the majority of non-syndromic CS remains unknown. Given that at least 79 genes are associated with CS, this suggests that multiple mechanisms can result in a similar outcome and leads us to propose that many of the genes converge on a common biological process that preserves suture patency.

We recently identified that during baso-apical calvarial growth, the ECM protein, Fibronectin 1 (FN1), was expressed in a graded manner towards the apex and enriched in the cranial mesenchyme apical of the calvarial primordia (Angelozzi et al., 2022; Feng et al., 2024). FN1 is a major component of the ECM in embryonic development involved in a wide range of contexts such as morphogenesis, cell adhesion, and migration (Danen and Yamada, 2001; Longstreth and Wang, 2024; Walma and Yamada, 2020; Walma and Yamada, 2022). *Fn1* deletion in the cranial mesenchyme derived from the neural crest and mesoderm preceding calvarial bone growth causes a decrease in apical expansion of the frontal bone and synostosis of the coronal suture (Feng et al., 2024). Interestingly, FN1 and other ECM proteins are upregulated in human periosteal fibroblasts of synostosed sutures from Apert syndrome, resulting from activating mutations in FGFR2 (Bodo et al., 1997; Lemonnier et al., 2001; Lomri et al., 1998). Conversely, in human Crouzon syndrome, which is also caused by activating mutations of FGFR2, FN1 protein expression is lower in suture-derived cells than controls (Baroni et al., 2002). From these findings, we hypothesize that dysregulation of FN1 expression is a downstream and shared etiology in CS, leading to disruption of the cranial suture patency.

In this study, we investigated cellular changes underlying coronal suture fusion in the *Fn1* mouse mutant and whether FN1 protein expression is dysregulated across mouse models of syndromic and non-syndromic CS. We found that conditional deletion of *Fn1* in the embryonic cranial mesenchyme causes ectopic osteogenic induction in the suture mesenchyme and subsequent fusion of the suture. Across different CS mouse models, graded FN1 protein expression along the baso-apical axis is dysregulated in the cranial mesenchyme with diminished apical expansion of the frontal bone primordia. These results highlight the crucial role of FN1 in sustaining cranial suture integrity and calvarial bone growth, shedding a new light on the mechanisms of calvarial development and the pathology of congenital calvarial defects such as CS.

## Methods

### Mouse strains and husbandry

*PDGFRα*‐*CreER* (Jax stock#018280) (Rivers et al., 2008), *Fn1* ^*flox/flox*^ (*Fn*^*fl/fl*^) (Jax stock #029624) (Sakai et al., 2001), Crouzon syndrome *Fgfr2c*^*C342Y/+*^ (Eswarakumar et al., 2004), Saethre-Chotzen syndrome *Twist1*^*+/-*^ (Chen and Behringer, 1995), Beare-Stevenson syndrome *Fgfr2*^*+/Y394C*^ (Wang et al., 2012) and *EIIA-Cre* ((Lakso et al., 1996) *Axin2*^*lacZ*^ (Jax stock# 009120) (Yu et al., 2005), *Tcf12*^*-/-*^ (Sharma et al., 2013) lines were used to study the role of fibronectin in calvaria and suture development. All mice were genotyped as described previously. For timed matings, *PDGFRα*‐*CreER/+; Fn*^*fl/fl*^ males were crossed with *Fn*^*fl/fl*^ females, and *Fgfr2*^*+/Y394C*^ males were bred with homozygous *EIIA-Cre* females. *PDGFRα*‐*CreER/+; Fn*^*fl/fl*^ mice were maintained on a mixed genetic background. The *Twist1*^*+/-*^, *Fgfr2*^*+/Y394C*^, and *EIIA-Cre* mice were maintained on the inbred C57BL/6J background. The *Fgfr2c*^*C342Y/+*^ mice were maintained on the outbred CD1 background. To yield the desired crosses, the mice were time mated overnight and checked for vaginal plugs in the morning. The vaginal plug day was assigned as embryonic day 0.5 (E0.5). The pregnant dams were given 25μg Tamoxifen/g body weight at E9.5 and 12.5μg Tamoxifen/g body weight at E10.5 by oral gavage to induce CreER recombination. For the knockdown model, mice were given 0.25μg Tamoxifen/g body weight at E9.5 and E10.5 to induce CreER recombination. Tamoxifen free base (Sigma, T5648) was dissolved in corn oil at 37℃ in a rotating chamber and used for oral gavage. Embryos were collected and processed as previously described(Atit et al., 2006). For each experiment, a minimum of four mutants with litter-matched controls from at least three litters were studied unless otherwise noted. Case Western Reserve University and Mount Sinai Institutional Animal Care and Use Committee approved all animal procedures in accordance with AVMA guidelines (Case Western: Protocol 2013–0156, Animal Welfare Assurance No. A3145–01; Mount Sinai: IPROTO202100000041).

### Histological stains, Immunohistochemistry, and Immunofluorescence

E13.5-E15.5 tissues were collected and fixed in 4% paraformaldehyde (PFA) at 4℃ for 35-50 min, respectively, sucrose dehydrated, embedded in O.C.T Compound (Tissue-Tek, Sakura Finetek USA, Torrance, CA, USA) and cryopreserved at -80℃. The embryos were cryo-sectioned at 14μm in the coronal or transverse plane. Immunofluorescence (IF) on cryosections was performed by drying slides at room temperature, washing in 1X PBS, and blocking in 5-10% donkey or goat serum. Sections were incubated with primary antibodies overnight at 4℃, washed in 1X PBS, incubated with species-specific secondary antibodies for 1 hour at room temperature, washed with DAPI (0.5 μg/mL), and then mounted with Fluoroshield (Sigma F6182). The following primary antibodies were used: recombinant rabbit anti-Fibronectin antibody (1: 250; Abcam ab26802; RRID:AB_2941028), rabbit anti-OSX (1:2000 or 1:4000; Abcam ab209484; RRID:AB_2892207), Goat anti-RUNX2 (1:100; R&D Systems AF2006-SP; RRID:AB_2184528), polyclonal rabbit anti-SIX2 (1:500; Proteintech 11562-1-AP; RRID:AB_2189084). Appropriate species-specific secondary antibodies included Alexa 488 (1:500, Invitrogen A32790; RRID: AB_2762833) and Alexa 594 (1:800 Invitrogen A11012; RRID: AB_2534079). OSX and SIX2 co-stain was performed by staining for each antigen sequentially. TUNEL assay was performed according to manufacturer directions (Thermo Fisher, C16017).

To quantify cell proliferation index, mice were administered 250ug EdU in PBS/10g of body weight by intraperitoneal injection 30 min. prior to sacrifice. Embryos were then collected, cryopreserved, and sectioned as described above. EdU was detected using Click-iT EdU Alexa Fluor 594 Imaging kit as per manufacturer directions (Invitrogen C10639). The slides were then stained with 1: 2000 DAPI for nuclei and mounted with Fluoroshield (Sigma F6182). The percent of proliferating cells OSX+ osteogenic fronts and coronal suture in the defined region of interest were manually quantified using Adobe Photoshop.

Alizarin red staining of cryosections was performed on sections after post-fixing in 4% PFA, washing in 1xPBS for 5 min, quickly rinsing with ddH_2_O before incubating with 2% Alizarin-S staining solution (Sigma Aldrich, TMS-008-C) for 15 seconds. The slides were washed with ddH_2_O and mounted with Vectashield mounting medium (H-1400-10). Alkaline phosphatase staining was performed on cryosections and apical extension of frontal bone primordia was quantified as previously described (Feng et al., 2024). Slides were scanned using Hamamatsu Nanozoomer S60 by 20X brightfield scanning microscope.

### Imaging

The brightfield images for histological stains were captured using the Olympus BX60 microscope with a digital camera (Olympus, DP70) and CellSens Entry Software (Olympus Corporation 2011; Version 1.5) with a 10X or 20X objective (Olympus UPIanFI 4/0.13). White balance was performed before the images were captured. The immunofluorescence images were captured on the Olympus BX60 wide-field microscope with Olympus DP70 camera. The exposure was held constant between controls and the experimental groups. Wholemount skeletal preparations were imaged with Leica MZ16F stereoscope and Leica DFC490 camera with Leica software. Images were processed using Adobe Photoshop and Fiji/ImageJ and page set in Adobe InDesign.

### RT-qPCR

The coronal suture on both sides of E15.5 control was micro dissected from control and CM*-Fn*^*fl/fl*^ embryos and RNA was isolated as previously described (Farmer et al., 2021; Ferguson et al., 2018). Relative mRNA expression was quantified using 4 ng of cDNA on a Step One Plus Real-Time PCR System (Life Technologies) and the ΔΔCT method (Schmittgen and Livak, 2008). Commercially available TaqMan probes (Life Technologies) specific to each gene were used: β*-actin* (ActB, 4352933E), *Six2* (Mm03003557_s1), *Erg* (Mm01214244_m1), *Efna2* (Mm00433011_m1), *Efna4* (Mm00433013_m1), *Twist1* (Mm00442036_m1). The CT values were normalized to β*-actin* CT value. ΔΔCT values were obtained by normalizing the ΔCT values to the average ΔCT values of the controls. Relative mRNA fold change was determined using the ΔΔCT values.

### Whole-mount skeletal preparation and Alizarin Red staining

For Alizarin Red whole-mount staining, embryos at E16.5 were fixed in 95% ethanol and acetone at 4℃ overnight before staining as previously described (Ferguson et al., 2018). Briefly, the embryos were pre-cleared in 1% potassium hydroxide (KOH) (Fisher Scientific, 1310–58-3) solution for 1 hour and stained in 0.005% alizarin red (Sigma, A5533) dissolved in 1% KOH overnight at 4℃. Then the embryos were placed in 50% glycerol (Sigma Aldrich, F6428-40ML): 1% KOH until clear. The embryos were kept in 100% glycerol for long-term storage. Images of controls and mutants were obtained at the same magnification on a Leica MZ16F stereoscope.

### Quantification and histomorphometry

At least 3 sections per embryo were used for histomorphometry. Quantifying the percent of cells expressing markers such as OSX and EdU was measured as the ratio of marker expressing number of cells to total DAPI stained nuclei number in the region of interest (ROI) boxes. ROIs were centered in the coronal suture between the frontal and parietal osteogenic fronts and were created to contain approximately 5 cell diameters of each osteogenic front in the control. Corrected fluorescence intensity was calculated from immunofluorescence images with Fiji/ImageJ. For fluorescence intensity (such as FN1), images were transformed to 16-bit and average of mean gray values of three equivalent circles on non-fluorescent regions was calculated as the background fluorescence using standard methods (Shihan et al., 2021). The corrected fluorescence per ROI was calculated by the following formula: Corrected fluorescence=Integrated Density-(Area of ROI×Average of background fluorescence). The line profile analysis was performed by Fiji/ImageJ software (Fiji/ImageJ; 2.14.0/1.54f). A Gaussian blur filter with a proper sigma (radius) value (15-20 in Fig 4, 5) was applied to the images. Four ROI lines were drawn from the basal to apical of the cranial mesenchyme on every image. Then the result CSV files were imported into a custom MATLAB script to obtain the intensity line profiles of the FN1 expression. The custom codes are available at http://github.com/radhikaatit/Apical expansion of calvarial osteoblasts and suture patency is dependent on fibronectin cues. The standard deviations were also incorporated into the plots as colored shaded area around the averaged intensity profiles.

### Statistics

All graphs and statistical analysis were generated using Microsoft^®^ Excel version 16.16.27 for Mac (Microsoft) and GraphPad Prism version 9.0.0 for Mac (GraphPad Software). Data are presented as mean ± SD in all graphs. Outliers were excluded using the outlier calculator in Microsoft^®^ Excel. For comparison of FN1 corrective fluorescence expression across multiple groups, quantification using ordinary One-way ANOVA multiple comparisons (Fisher’s). For comparisons between two groups, unpaired, two-tailed Welch’s *t*-test was performed due to unequal variance and sample numbers. The p-values for statistical tests in all figures are represented as: *=p<0.05, **=p<0.001, ***=p<0.0001 and ****=p<0.00001.

## Results

Previous studies report the dysregulation of FN1 in human calvarial suture fibroblasts from CS patients(Baroni et al., 2002; Bodo et al., 1997). In the mouse embryo between E12.5-E13.5, we reported baso-apically graded expression of FN1 expression in the cranial mesenchyme that included progenitors for the dermis and frontal bone primordia (Feng et al., 2024). Our prior work showed that deletion of FN1 expression in the cranial mesenchyme before calvarial apical expansion was sufficient to cause CS in mouse embryos; however, the underlying mechanism for this effect remains to be elucidated.

### Loss of FN in cranial mesenchyme causes CS in coronal suture

To understand the role of FN1 in suture patency, we used a tamoxifen-inducible conditional deletion of *Fn1* in the cranial mesenchyme starting at E9.5 using *Pdgfrα-CreER*-mediated recombination as previously described and validated in (Feng et al., 2024) (CM-*Fn*^*fl/fl*^; **Figure 1A**). Using immunostaining, we found expression of FN1 protein in the transverse plane is visible in the coronal suture, osteogenic fronts of the frontal and parietal bone primordia, and in the underlying meninges and in the overlying dermis. In E13.5 CM-*Fn*^*fl/fl*^ mutants, FN1 protein expression was nearly undetectable in coronal suture, osteogenic fronts of the frontal and parietal bone primordia (**Figure 1B, C, B’, C’**). By E15.5 -E16.5, we found ectopic mineralized bone in the coronal suture of CM-*Fn*^*fl/fl*^ embryos (**Figure 1E-G**). CM-*Fn*^*fl/fl*^ embryos are embryonically lethal precluding postnatal analysis These results raised the question of whether loss of FN1 causes loss of patency by disrupting formation or maintenance of the coronal suture mesenchyme.

**Figure 1.**
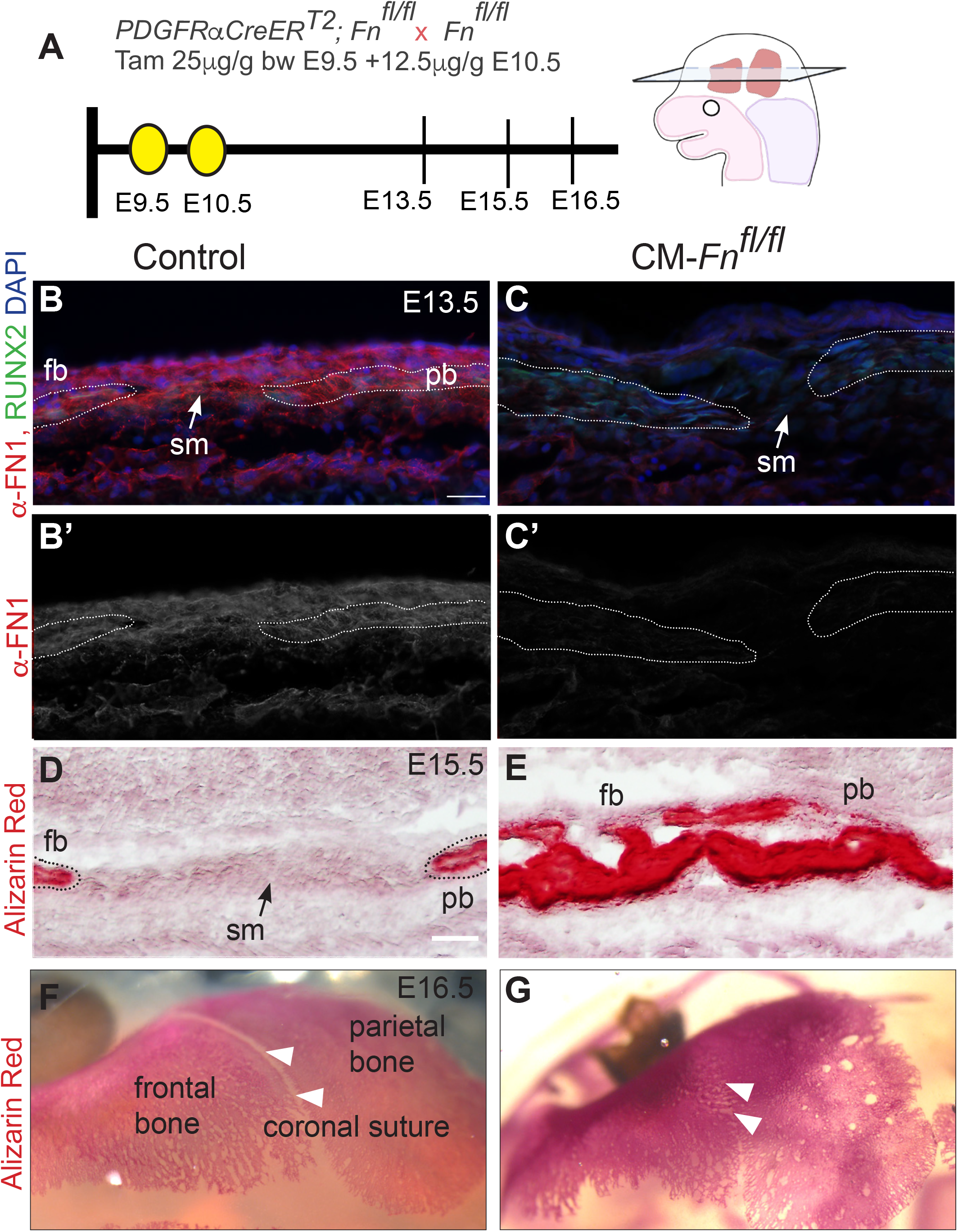
Inducible model of FN1 gradient loss leading to coronal suture fusion. (A) Schematic of tamoxifen induced deletion of *Fn1* in the cranial mesenchyme (CM-*Fn*^*fl/fl*^) and orientation of transverse sections. (B, B, C, C’) Indirect immunofluorescent staining for FN1 protein (in red) in the coronal suture of control and CM-*Fn*^*fl/fl*^ mutants. The frontal and parietal bone primordia are marked by RUNX2 expression (green) and outlined in dotted white lines. (D-E) Alizarin Red staining in sections black dotted lines mark the mineralization fronts. (F, G) Wholemount dorsal view showing mineralization in the coronal suture in CM-*Fn*^*fl/fl*^ mutants at the same magnification (white arrowheads in G). *fb*, frontal bone; *pb*, parietal bone; *sm*, coronal suture mesenchyme. Arrows and arrowheads mark the coronal suture. (B) Scale bar=25μM; (D) Scale bar =100μM. Schematics were made using biorender.com

### Suture mesenchyme survival and initial demarcation of the osteogenic and suture mesenchyme boundaries occurs independently of Fibronectin function

Suture mesenchyme in the coronal suture is specified by E12.5 and expands baso-apically with calvarial primordia in the mouse embryo (Deckelbaum et al., 2012; White et al., 2021). Fusion of the coronal suture by E15.5-16.5 in CM-*Fn*^*fl/fl*^ mutants could occur by several means of altered cell behavior. One plausible mechanism is that suture patency could be lost due to defects in suture mesenchyme survival and thus premature fusion. Analysis of cell death by TUNEL assay did not reveal apoptotic cells in the coronal suture mesenchyme cells in E13.5 controls or CM-*Fn*^*fl/fl*^ mutants suggesting that this mechanism was not prevalent (**Supplementary Figure 1A-B’)**.

We next assess if FN1 function led to inappropriate cell dynamics and integration at osteogenic fronts of the frontal and parietal bone. Genetic lineage analysis in mouse embryos shows the coronal suture marks the boundary between the neural crest-derived frontal bone and the paraxial mesoderm-derived parietal bone and suture mesenchyme (Jiang et al., 2002; Yoshida et al., 2008). In *Twist1*^*+/-*^ and *Epha4* ^*-/-*^ embryos, there is mixing of neural crest and mesoderm at the coronal suture lineage boundary (Merrill et al., 2006; Ting et al., 2009). *Twist1*^*+/-*^ coronal sutures have reduced expression of *Ephrin A2* (*Efna2)* and *Ephrin A4* (*Efna4)*, and *Epha4* is a functional downstream effector of *Twist1* in CS of the coronal suture (Merrill et al., 2006; Ting et al., 2009). However, in contrast, lineage mixing was not observed in the *Fgfr2*^*+/S252W*^ Apert syndrome model (Holmes and Basilico, 2012). To determine the etiology underlying the loss of coronal suture in CM-*Fn*^*fl/fl*^ mutants, we queried the mRNA expression at E15.5 in the coronal suture. We found that expression of *Epha2, Epha4* and *Twist1* mRNA expression was comparable between controls and CM-*Fn*^*fl/fl*^ mutants, suggesting that the loss of coronal suture patency is also not likely due to disruption of the lineage boundary regulated by Twist1-Eph-Ephrin signaling (**Supplementary Figure 1C-F**).

### Fibronectin deficiency leads to precocious osteogenic induction and a decrease in proliferation in the coronal suture

One hypothesis of the suture fusion resulting from FN1 deficiency is a shift in the differentiation state of existing cells. We investigated if FN1 is required to prevent osteogenic induction in the suture mesenchyme. We visualized the differentiation status and proliferation index of cells in the suture within the transverse plane of the skull at E14.5, 24 hours prior to morphologically detectable CS by mineralization of the coronal suture that occurs between E15.5-16.5 in the mutant. The CM-*Fn*^*fl/fl*^ mutant phenotype varied from a reduced suture to the complete loss of osteogenic/non-osteogenic suture boundary integrity in the apex of the skull; a representative region is shown in **Figure 2A, B**. Compared to controls, we found that osteogenic fronts of the frontal and parietal bone primordia in the transverse plane in the coronal suture were qualitatively thicker and less tapered CM-*Fn*^*fl/fl*^ mutant (**Figure 2A-B”**). We quantified the proliferation index of cells in the region of interest (ROI) that includes presumptive coronal suture and osteogenic front tips as previously described (Teng et al., 2018). Proliferation was comparable between mutants and WT in the ROI (**Figure 2C)**. However, relative to controls, CM-*Fn*^*fl/fl*^ mutants had an increased percentage of OSX^+^ osteogenic cells, presumably due to the expression of OSX throughout the suture mesenchyme, which was not present in controls (**Figure 2D**). The percentage of proliferative OSX+ calvarial osteoblasts was significantly less in the CM-*Fn*^*fl/fl*^ mutant (**Figure 2E**). These data suggest that cells in the suture of CM-*Fn*^*fl/fl*^ mutants are differentiating towards an osteogenic program at the expense of proliferation. Together, these findings suggest that FN1 preserves coronal suture patency by suppressing ectopic differentiation.

**Figure 2:**
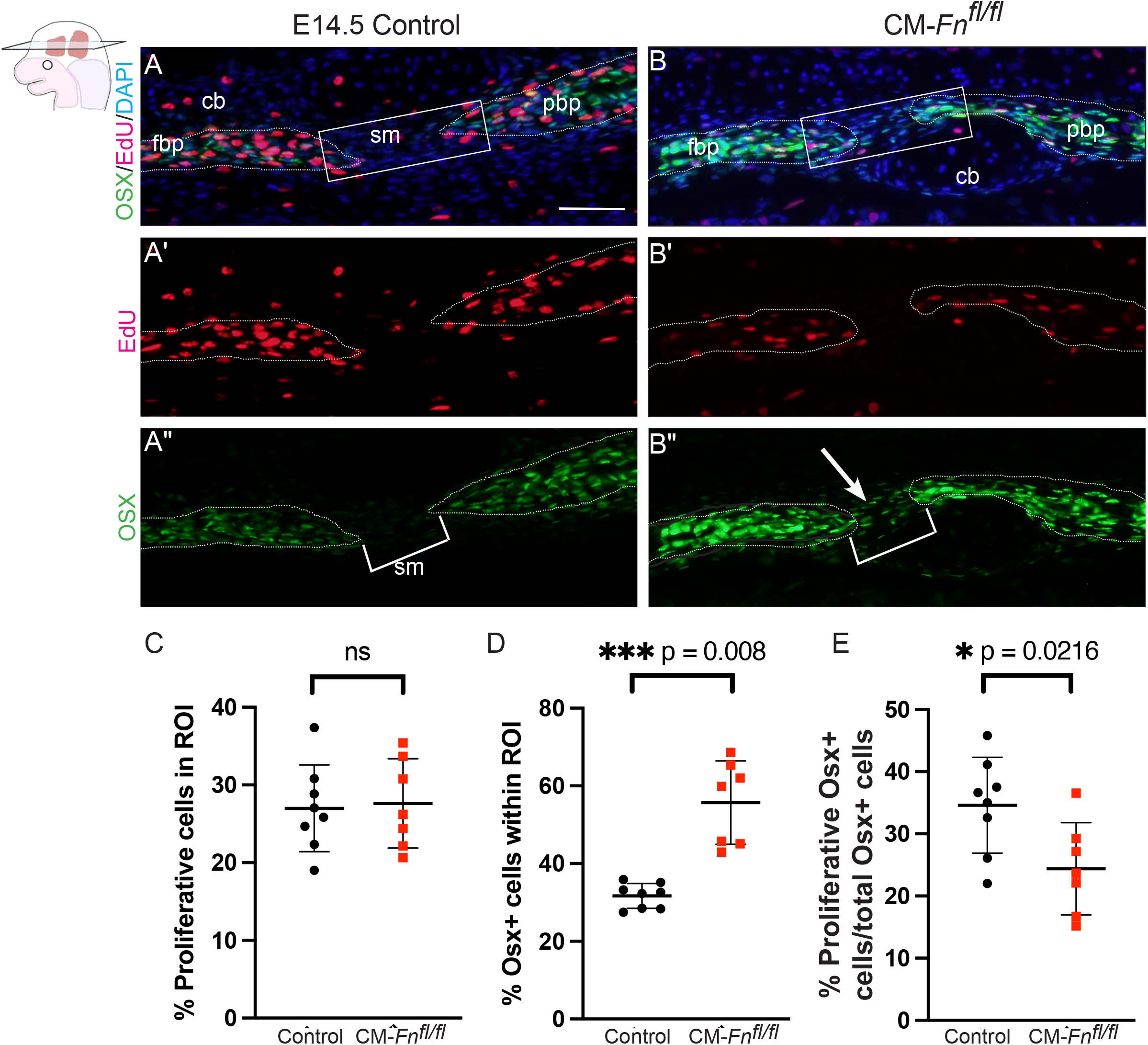
Precocious osteogenic differentiation in the suture mesenchyme with loss of fibronectin function. (A, B) Protein expression of OSX (green) by indirect immunofluorescence and EdU incorporation (red) in E14.5 transverse sections through the coronal suture between the frontal bone primordia (fpb) and parietal bone primordia (pbp). Region of interest (ROI) is demarcated (white box) (C) Proliferation index of cells in the ROI in controls and CM-*Fn*^*fl/fl*^ mutants. (D) Percentage of OSX^+^ cells in the ROI in the coronal suture of CM-*Fn*^*fl/fl*^ mutants is significantly increased relative to the control. (E) Percentage of proliferative OSX^+^ cells is significantly reduced in the coronal suture of CM-*Fn*^*fl/fl*^ mutants (8 controls and 7 mutants). Data are means +/-s.d.; *ns*, not significant *p*≥0.05, \**p*≤0.05, \*\*\**p*≤0.0005 (Welch’s two-tailed *t*-test). Abbreviations: *sm*, suture mesenchyme; *cb*, cartilage base; Scale bar=50μM. Schematics were made using biorender.com

Next, we investigated if coronal suture mesenchyme identity was compromised or co-existed with ectopic osteogenic induction in CM-*Fn*^*fl/fl*^ mutants. Recent single-cell RNA sequencing of the coronal suture identified marker genes such as transcription factors *SIX homeobox 2* (*Six2*) and *Erg* that are enriched in the E15.5-18.5 coronal suture (Farmer et al., 2021; Holmes et al., 2021). Compared to E15.5 controls, in CM-*Fn*^*fl/fl*^ mutants we found *Six2* mRNA to be slightly lessened while *Erg* mRNA to be significantly decreased in coronal suture enriched mesenchyme by quantitative RT-PCR(**Figure 3A-C**), suggesting a broad decrease, or loss, of suture mesenchyme markers within the population of suture cells from the CM-*Fn*^*fl/fl*^ mutant. To obtain spatial information, we investigated protein expression of SIX2 and OSX during calvarial suture mesenchyme specification and osteogenic induction. In the E15.5 presumptive coronal suture of CM-*Fn*^*fl/fl*^ mutants, the osteogenic fronts were not clearly defined, SIX2 expression was absent, and ectopic osteogenesis appeared to have replaced the suture. The earliest detectable protein expression of SIX2 in the suture mesenchyme cells was at E14.5 (**Figure 3 D)**. In wildtype controls, we did not detect co-expression of SIX2 and OSX in cells in the coronal suture (n=4). In E14.5 CM-*Fn*^*fl/fl*^ mutants, SIX2 protein expression was visible in the coronal suture suggesting the suture mesenchyme is specified (**Figure 3D, E)**. In contrast to controls, we found co-expression of OSX and SIX2 in the center region of the suture in E14.5 CM-*Fn*^*fl/fl*^ mutants (n=4/4) (**Figure 3F**). Furthermore, the suture architecture was altered in CM-*Fn*^*fl/fl*^ mutants with SIX2^+^ cells more dispersed around the thickened osteogenic fronts (n=4/4) (**Figure 3E**). These results together suggest that SIX2^+^ suture mesenchyme is specified and visible in the patent regions of the coronal suture in CM-*Fn*^*fl/fl*^ mutants at E14.5, but the ectopic osteogenic program is emerging in the SIX2^+^ cells, showing that FN1 is required to maintain suture mesenchyme identity and patency while suppressing the osteogenic program (**Figure 3G**).

**Figure 3.**
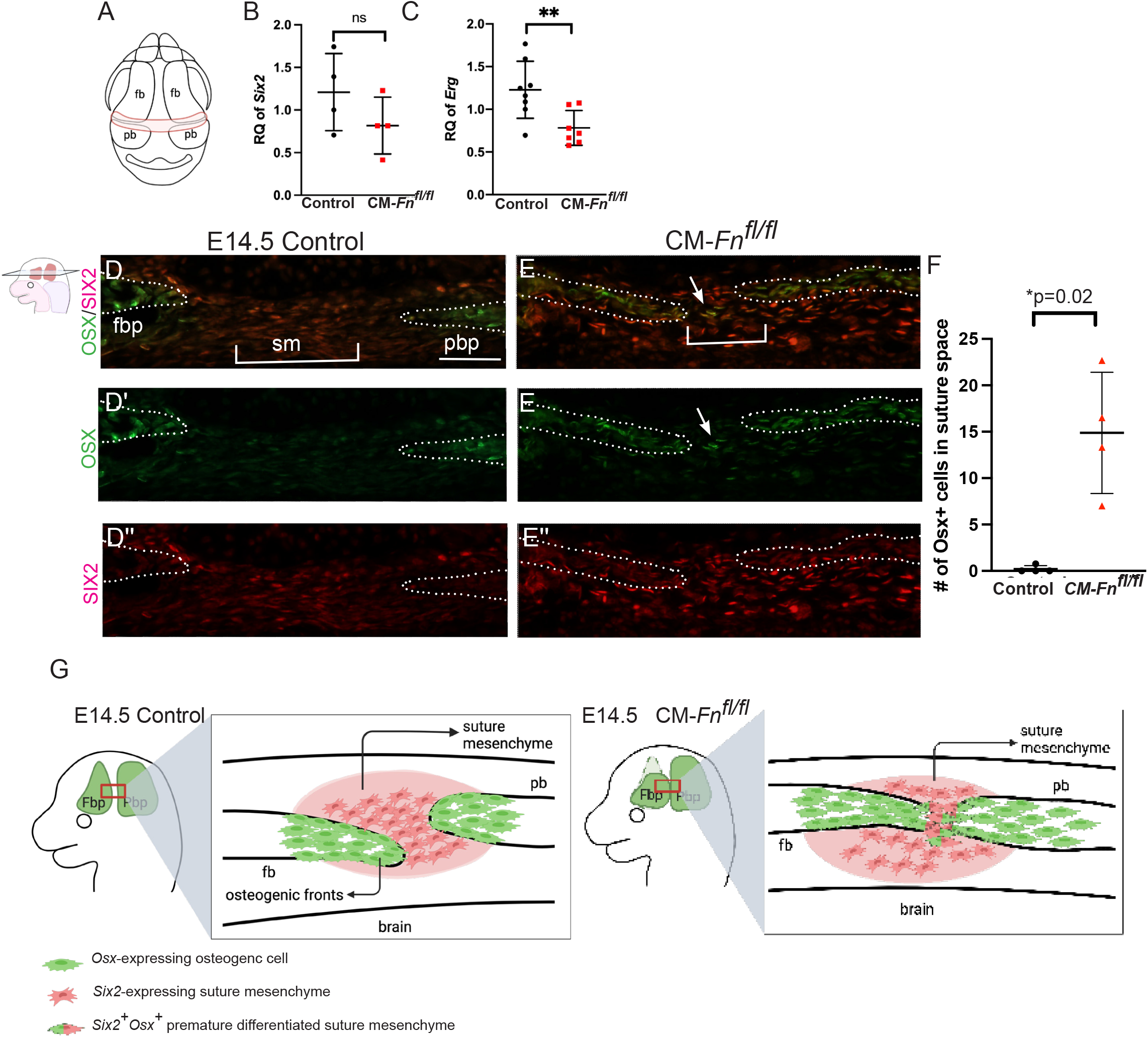
Loss of Fn1 induces premature osteogenesis in coronal suture. (A) Schematic depicting the tissue source of RT-qPCR analysis of E15.5 coronal sutures (B, C) RT-qPCR analyses showing relative mRNA quantities of *Six2* and *Erg* mRNA in controls and CM-*Fn*^*fl/fl*^ mutants at E15.5 (n= 4, controls; 4 mutants). (D, E) Transverse section through the coronal suture with indirect immunofluorescent staining for SIX2^+^ (red) suture mesenchyme and OSX^+^ (green) osteogenic fronts. (D’,D”, E’, E”) Compared to controls, some of the suture mesenchyme cells in CM-*Fn*^*fl/fl*^ mutants have co-expression of SIX2 and OSX (white arrow) in the coronal suture mesenchyme and SIX2 expression domain is expanded around the osteogenic fronts (F) Number of OSX+ cells in the center of the suture/fixed ROI in controls and mutants (n=4 controls and mutants) (G) A schematic of the working model illustrating that ectopic osteogenesis in the suture mesenchyme of CM-*Fn*^*fl/fl*^ mutants at E14.5. *fbp* frontal bone primordia; *pbp*, parietal bone primordia, *sm*, suture mesenchyme of the coronal suture. Scale bar=100μM. \**p*≥0.05 (not significant), \*\**p*≤0.005, (Welch’s *t*-test). Schematics were made using biorender.com

### FN1 graded expression and resulting apical expansion is disrupted across mouse models of CS stemming from different genetic causes

FN1 expression is dysregulated in human CS suture fibroblasts *in vitro* (Baroni et al., 2002; Bodo et al., 1997). We have also shown the FN1 protein expression is elevated in the cranial mesenchyme of *Fgfr2*^*+/S252W*^ Apert mouse model during calvarial development (Feng et al., 2024). Given the common pathology of suture fusion in CS stems from different genetic and environmental causes, we queried if a common mechanism of FN1-mediated calvarial apical expansion may be affected across cases of CS originating from different genetic contexts. As this line of inquiry cannot be investigated in patient samples, we investigated different, validated, mouse models of CS to ask if dysregulation of graded FN1 protein expression and apical expansion of calvaria in embryonic development is shared among syndromic and non-syndromic CS models.

We visualized and quantified FN1 protein expression in embryonic cranial mesenchyme in coronal plane sections using line graph analysis and corrected fluorescence in five different mouse models of CS (**Figures 4 & 5**). In E13.5 controls, FN1 protein expression is graded along the baso-apical axis of the cranial mesenchyme as seen by the line graphs. In the Crouzon syndrome model with a *Fgfr2c*^*C342Y/+*^ activating mutation, we found the graded expression of FN1 was consistently diminished compared to the control (**Figure 4A-C, H)**. The apical extension of the frontal bone was also significantly diminished at E13.5 (**Figure 4L**). Similarly, we found diminished FN1 protein expression in the cranial mesenchyme and decreased apical expansion of the frontal bone primordia in the Beare-Stevenson syndrome CS model with a *Fgfr2*^*Y394C/+*^ mutation(Wang et al., 2012) (**Figure 4I, M Supplementary Figure 2)**. In the *Twist1*^*+/-*^ heterozygous mouse mutant, a model for Saethre-Chotzen syndrome, FN1 protein expression was markedly elevated throughout the cranial mesenchyme and there was loss of graded FN1 expression along the baso-apical axis as seen by line graph and corrective fluorescence (**Figure 4 D-E, J)** (Behr et al., 2011; Chen and Behringer, 1995). The normalized apical expansion of frontal bone was significantly diminished in *Twist1*^*+/-*^ mutants (**Figure 4N**). The *Axin-2*^*LacZ/ LacZ*^ mutant is a non-syndromic model and develops CS postnatally (Yu et al., 2005). In the *Axin2* ^*LacZ/ LacZ*^ embryos, we found significantly elevated expression of FN1 protein in the apical region and accompanied by significantly diminished expansion of the frontal bone primordia (**Figure 5A-C, H, K)**. Similarly, in another non-syndromic mouse model with *Tcf12*^*-/-*^ null mutation, we found elevated expression of FN1 protein expression (**Figure 5 D-F, I, L)** (Sharma et al., 2013; Twigg and Wilkie, 2015). Thus, dysregulation of graded FN1 expression accompanied by impaired apical expansion defects in frontal bone primordia was evident across five different CS mouse models of disparate genetic etiologies and lends support to the idea that there is a convergence point and shared etiological mechanism (**Figure 6**).

**Figure 4.**
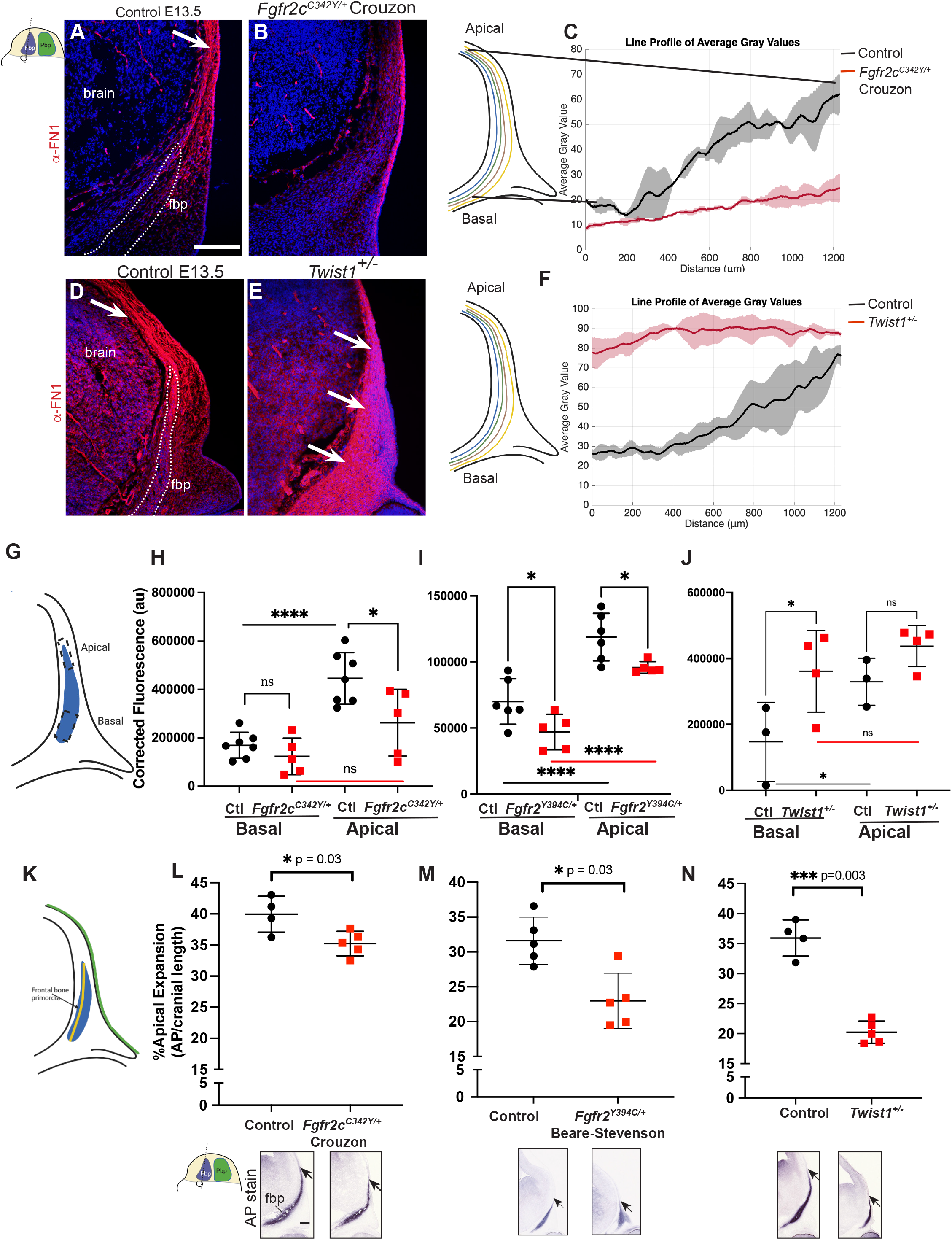
*Fgfr2c*^*C342Y/+*^, *Fgfr2*^*Y394C*^*/+*, and *Twist1*^+/-^ mutations leads to loss of graded FN1 expression that associates with decreased apical expansion of the frontal bone primordia. (A, B, D, E) Representative images showing immunofluorescence of FN1 in E13.5 coronal sections revealing altered FN1 expression in various mutant embryo relative to control. (C, F, G-J) Quantification of FN1 expression by line graph and corrective fluorescence in cranial mesenchyme showing changes in graded FN1 expression compared with control. In C, F, average gray value is shown as the lines, and the standard deviation of the quantifications is shown as colored shades. (K) Schematic for quantifying normalized percent apical extension of AP^+^ frontal bone primordia. (L-N) AP staining in E13.5 coronal sections showing decreased apical extension of frontal bone primordia in different mutants. Each value is the average of 3-4 sections from one embryo. N=3-8 controls; 3-5 mutants. In H-J, L-N data are mean ± s.d. p≥0.05 (not significant; ns), *p<0.05, ***p≤0.0005 with One-Way ANOVA (H-J) and Welch’s *t*-test (L-N). fbp, frontal bone primordia. Scale bar=100μM. Schematics were made using biorender.com

**Figure 5:**
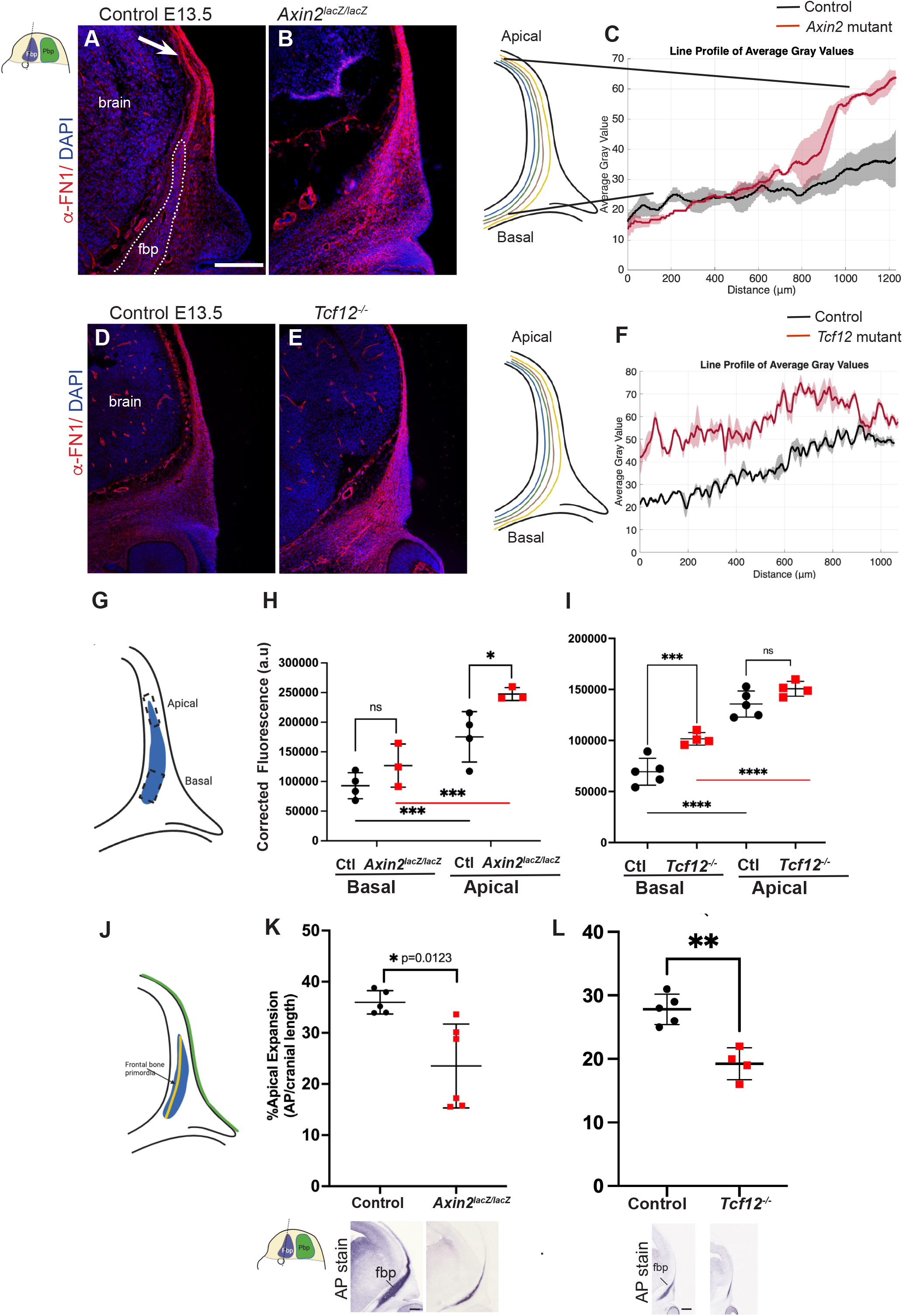
In *Axin2*^*lacZ/lacZ*^ and *Tcf12*^*-/-*^ non-syndromic CS mutants, FN1 expression is elevated, and the apical expansion of frontal bone primordia is significantly reduced. (A, B, D, E) Representative images showing immunofluorescence of FN1 in E13.5 coronal sections revealing increased FN1 in *Axin2*^*lacZ/lacZ*^ and *Tcf12*^*-/-*^ mutant. (C, F, G-I) Quantification of FN1 expression by line graph and corrective fluorescence in cranial mesenchyme shows the *Axin2*^*lacZ/lacZ*^ and *Tcf12*^*-/-*^ mutants. N=4 controls; 3 litter-matched mutants for corrected fluorescence quantification. In C and F, average gray value is shown as the lines, and the standard deviation of the quantifications is shown as colored shades. (F, G, I) (J) Schematic for quantifying normalized percent apical extension of AP^+^ frontal bone primordia. (K, L) AP staining in E13.5 coronal sections showing decrease in normalized apical length of frontal bone primordia. N=4-5 controls; 5-6 mutants. In H, I, K, L data are mean ± s.d. P≥0.05 (not significant; ns), *p<0.05, ** p≤0.005, ***p≤0.0005 with One-Way ANOVA (H, I) and Welch’s *t*-test (K, L). Scale bar=100μM. Schematics were made using biorender.com

**Figure 6:**
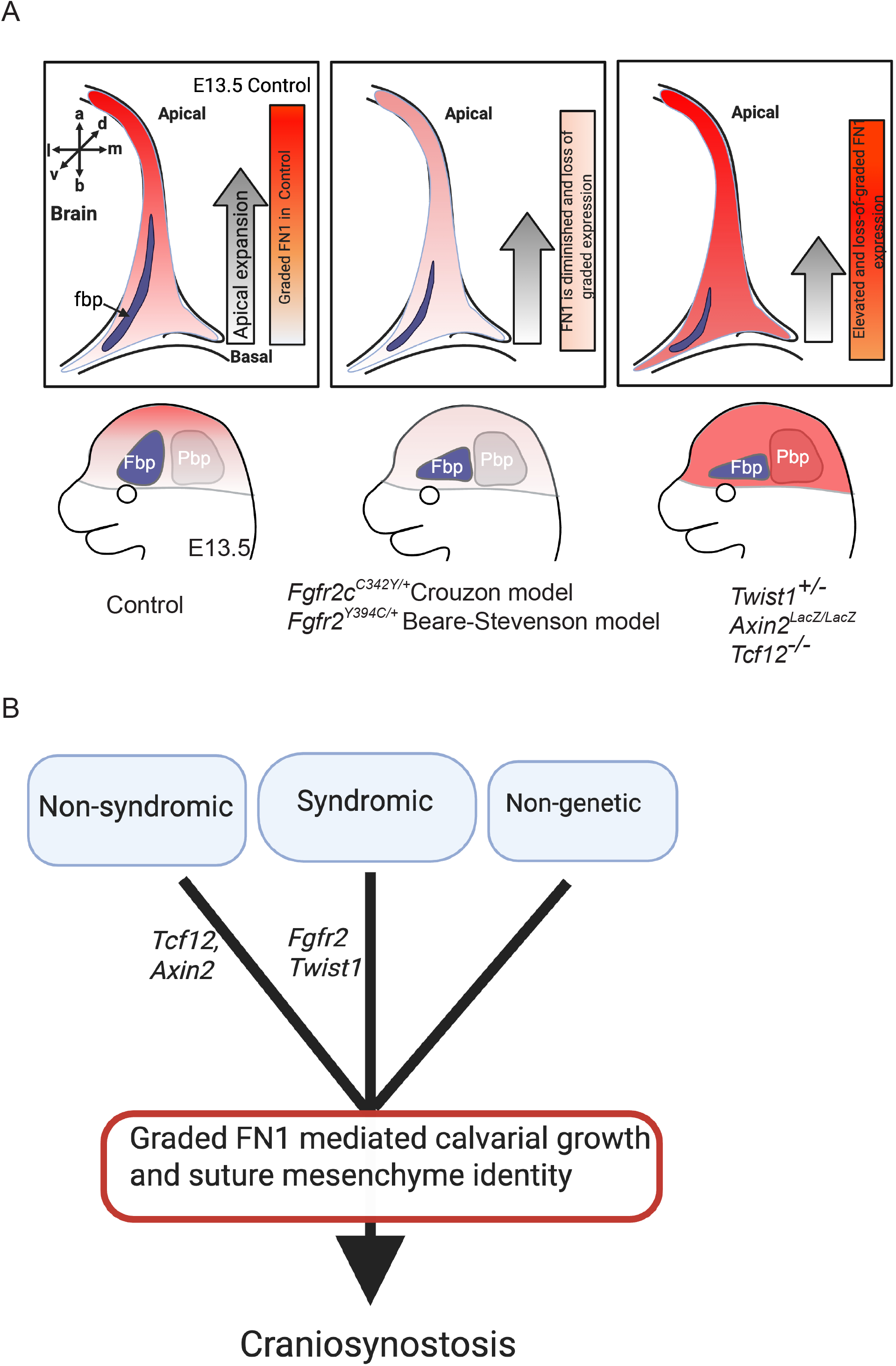
Summary of dysregulated FN1 expression in the cranial mesenchyme and apical expansion in 5 different CS models of different etiologies and working model. (A) Summary schematic showing dysregulated FN1 expression in CM of 5 different CS mouse models accompanied with diminished apical expansion of the frontal bone primordia. fbp, frontal bone primordia. (B) Working model suggesting that mutation in genes associated with CS leads to disruption of a common biological process that leads to loss sutures in CS. Schematics were made using biorender.com. https://BioRender.com/ztprfcq

## Discussion

Although numerous genetic causes of CS have been identified, we remain without a central etiology on how the dysregulation of these factors lead to altered calvarial growth and disruptive suture formation. Previously, we identified the extracellular matrix as a possible integrating component of CS, based on our identification of a FN1 expression gradient that is essential for apical calvarial expansion and suture patency during development. Disruption of the FN1 gradient is associated with Apert model of CS (Feng et al., 2024). Here, we build on these findings to define how FN1 mediates suture patency, and assess changes in calvarial growth and FN1 gradient across different mouse models of CS. We found that loss of *Fn1* in CM does not seem to affect cell viability nor loss of identity but instead promotes premature differentiation of the coronal suture mesenchymal cells to osteoblasts. Moreover, disruption of the FN1 gradient –whether by decreased or enhanced expression – is consistently associated with apical expansion of frontal bone in five distinct CS models. These findings are in concordance with variance of FN1 seen in suture fibroblasts from human CS patients (Al-Qattan et al., 2014; Baroni et al., 2002; Bodo et al., 1997). Collectively, our results support a model in which a spatially graded extra-cellular environment along the basal-to-apical axis of the developing skull coordinates normal apical expansion and suture formation. Loss of this directional polarity disrupts normal apical growth and contributes to premature suture fusion.

FN1 plays multiple essential roles in morphogenesis as a substrate for cell migration, extracellular matrix organization, and mechanical signaling (Chen et al., 2007; Chen et al., 2021; Danen and Yamada, 2001; Darribère and Schwarzbauer, 2000; Martino et al., 2018; Vega and Schwarzbauer, 2016; Yamada and Sixt, 2019; Yamada et al., 2022). FN1 is well known as an adhesive substrate for cell migration. Calvarial osteoblasts are migratory cells and depend on Arp2/3-WASL1-dependent lamellipodia for apical expansion of the frontal bone *in vivo* and *in vitro* (Feng et al., 2024; Liu et al., 2024). Consistent with this behavior, calvarial osteoblast precursors exhibit Wnt5a-dependent protrusions *in vivo*, further supporting a role for active cell-movement during skull morphogenesis (Liu et al., 2024; Polsani et al., 2024). We recently demonstrated that calvarial osteoblasts can migrate directionally along a graded concentration of FN1 *in vitro* (Liu et al., 2024). In line with this finding, we show here that dysregulation of FN1 expression across multiple CS models coincides with impaired baso-apical calvarial expansion. These results suggest that altering the graded expression of FN1 interferes with cell-ECM interactions or cell motility that contribute to pathological calvarial expansion. Beyond its role in migration, the fibrillar scaffolding features of FN1 may provide mechanical cues for apical expansion of the calvaria and for the elongation of the osteogenic fronts, thereby contributing to proper coronal suture architecture. Integrin binding to FN1 enables transmission of mechanical forces between cells and the ECM, a process likely important for osteoblasts movement through the cranial mesenchyme towards apex and for shaping the advancing osteogenic fronts (Danen and Yamada, 2001; Longstreth and Wang, 2024; Yamada et al., 2022). Consistent with this model, the osteogenic fronts of the frontal and parietal bone primordia in the coronal suture are thicker and less tapered in the CM-*Fn*^*fl/fl*^ mutants. Such alterations in suture and osteogenic front architecture may disrupt the paracrine signaling between osteogenic front and intervening suture mesenchyme cells, thereby contributing to loss of suture patency(Holmes and Basilico, 2012). In addition, FN1 is known to organize other ECM components, including collagen and laminin, and to bind and present growth factors (Danen and Yamada, 2001; Longstreth and Wang, 2024; Yamada et al., 2022). These functions may be required to generate an ECM environment that permits osteogenic fronts to elongate and expand while also preserving suture patency. Supporting this idea, recent studies ex-vivo studies suggest that collagen matrix assembly may be important in the organization of the osteogenic fronts in the metopic suture (Dang et al., 2025). Future studies that selectively perturb FN1-integrin interactions via altering the RGD domain, modulate FN1 stiffness, or examine growth factor signaling and transcriptional changes in the absence of FN1 will be essential to disentangle the distinct mechanical and signaling roles of FN1 in calvarial growth and suture patency.

In this study, we deleted FN1 broadly starting at E9.5 in the *PDGFRα-CreER*-derived cranial mesenchyme that gives rise to frontal and parietal bone, suture, meninges, and dermis of the skin (Ferguson et al., 2018), and see synostosis of the coronal suture either unilaterally or bilaterally by E15.5 which is relatively early for developing CS. Our previous study defined a window in which FN1 was required to preserve suture patency, and deletion of FN1 after E14.0 did not lead fusion of the coronal suture (Feng et al., 2024). Thus, the temporal role of FN1 in preserving suture patency before E14.0 may be a result in the timing of the FN1 deletion and alteration of growth factors that may be relevant in the coronal suture. Which cell type uses FN1 to preserve suture patency is unclear from our studies. It is not clear if it is a paracrine effect from neural crest or mesoderm derived lineages or cell-autonomous to the suture mesenchyme. It may also be altered juxtracrine signaling between osteogenic fronts and adjacent suture mesenchyme cells to induce osteogenesis. The developing mammalian coronal suture forms at the cranial neural crest and mesoderm lineage boundary and between osteoblasts and suture mesenchyme cell type boundary (Jiang et al., 2002; Yoshida et al., 2008). Loss of either of these boundaries can occur in mouse models of CS(Holmes and Basilico, 2012; Merrill et al., 2006). The relative importance of both these boundaries is unclear in the CM-*Fn*^*fl/fl*^ mutant, but our results suggest that *Twist1-Eph-Ephrin* mRNA expression is not altered preceding the loss of the coronal suture. We found the induction of ectopic osteogenesis in the suture mesenchyme in the CM-*Fn*^*fl/fl*^ mutant preceding the premature loss of the coronal suture. These results suggest that FN1 is required to promote or preserve the stemness of the suture mesenchyme and prevent the osteogenesis program. We see initial formation of suture mesenchyme, but the subsequent osteogenic induction within this mesenchyme suggests a later loss in cell type identity or cell-autonomous fate change to osteogenic lineage. As lineage-specific Cre lines that can delete FN1 after E9.5 in different cell types become available, we can test which cell type requires FN1 expression to preserve suture patency.

Understanding the interplays among ECM dysregulation, calvarial bone growth and CS holds significant clinical implications. It paves the way for developing targeted therapies that could modulate ECM components or the signaling pathways involved in their regulation. Currently, the primary treatment for CS involves surgical intervention to correct the skull deformity and prevent further complications. However, with advancements in genetic and molecular biology, future treatments may include gene therapy or molecular inhibitors that specifically address the underlying causes of ECM dysregulation. Continued research into the ECM’s role in CS further promises to refine therapeutic approaches and improve the treatment outcomes for affected individuals.

## Acknowledgements

We thank all the members of the Atit, Holmes, and Harris Lab past and present for helpful comments and suggestions in guiding this project. We thank Melissa Carr with help on animal handling. We thank Chao Liu for help with generating line graph analysis code. Thank you to Case Western Reserve University bio[box] facility for use of the shared instrumentation facility. All schematics were made on biorender.com.

## Competing interests

The authors declare no competing or financial interests

## Author Contributions

Conceptualization: X.F., R.P.A, G.H., M.P.H

Methodology: X.F, M.G., R.P.A, G.H., M.P.H

Software: X.F. Validation: X.F., R.P.A

Formal analysis: X.F. A.F. M.G., M.H., R.P.A

Investigation: X.F. A.F. M.G., M.H., R.P.A, G.H., D’J.F. M.S.

Resources: R.P.A, G.H., D’J.F. M.S.

Writing original draft: X.F., R.P.A, G.H.,

Writing-review & Editing: X.F., R.P.A, G.H., M.P.H

Visualization: X.F., R.P.A, G.H., M.P.H

Supervision: X.F., R.P.A, G.H.

Project administration: X.F., R.P.A, G.H.

Funding acquisition: R.P.A, G.H., M.P.H

## Funding

This work was supported by National Institutes of Health/National Institute of Dental and Craniofacial research (R56 DE030206, R01 DE032670 R.P.A, M.P.H, G.H.), Case Western Reserve University ENGAGE summer fellowship to A.F.

## Data availability

The custom MATLAB script for obtaining the intensity line profiles of the FN1expression are available at https://github.com/atit-lab/FN1-line-graph-analysis. All other relevant data can be found within the article and its supplementary information.

## Notes

### Competing Interest Statement

The authors have declared no competing interest.

