## supplementary figures for "Fibronectin 1 is required for suture patency and dysregulated across craniosynostosis models in the mouse"

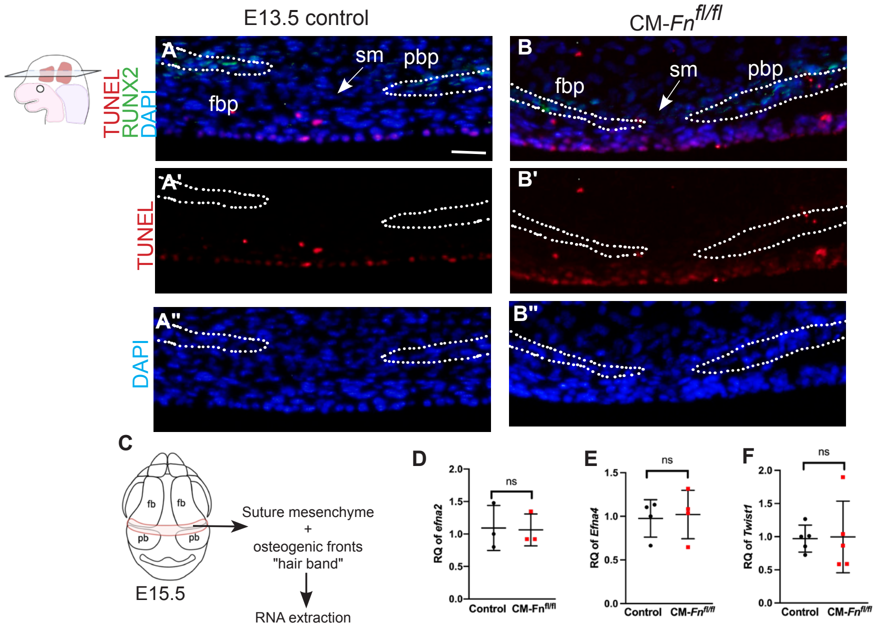


Feng et al.,
Supplementary Figure 1

**Supplementary Fig 1: Cell survival and expression of lineage boundary genes in suture mesenchyme cells is comparable between control and CM-Fn$^{fl/fl}$ mutants.** (A, B, A', B', A", B") Transverse sections of E13.5 controls and CM-Fn$^{fl/fl}$ mutants did not show overt cell death by TUNEL assay in the coronal suture (cs). Dotted lines outline the osteogenic fronts of frontal bone primordia (fbp) and parietal bones primordia (pbp). (C) Schematic depicting the tissue source and workflow for the RT-qPCR analysis of E15.5 coronal sutures. (D-F) RT-qPCR analyses showing relative mRNA quantities (RQ) of *Ephrin a2 (Efna2), Ephrin a4 (Efna4), Twist1* mRNA in controls and CM-Fn$^{fl/fl}$ mutants at E15.5. N=3-5 controls; 3-5 mutants. Data are means + s.d. ns, not significant p≥0.05, Welch's two-tailed t-test. Scale Bar=100μM.

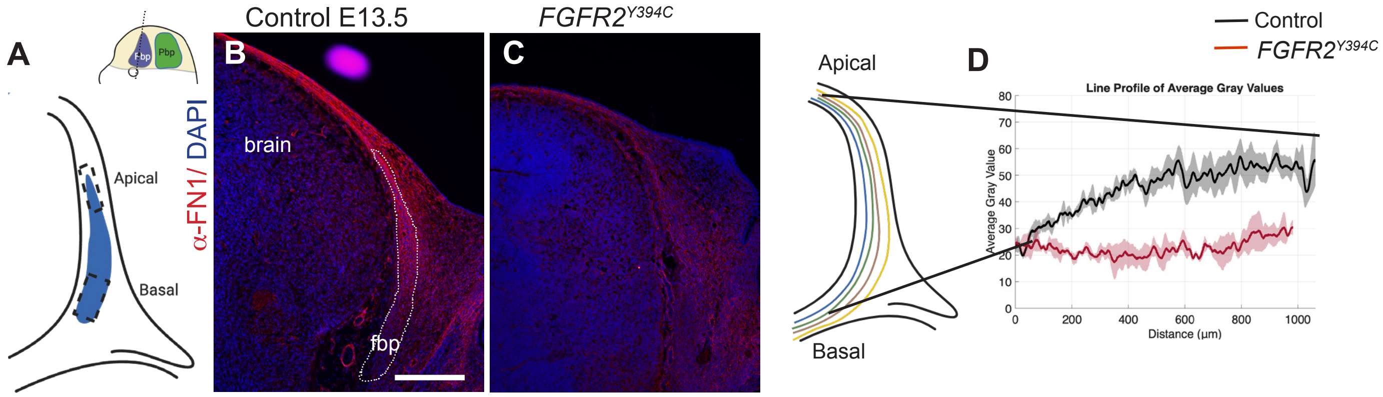


Feng et al.,
Supplementary Figure 2

**Supplementary FIgure2. Fgfr2$^{Y394c/+}$ mutation leads to loss of graded FN1 expression.** (A) Schematic of a coronal section with the frontal bone primordia (blue, fbp) (B,C) Representative images showing immunofluorescence of FN1 in the cranial mesenchyme at E13.5 in control revealing altered FN1 expression in mutant embryo relative to control. (D) Visualization of FN1 expression in cranial mesenchyme showing changes in graded FN1 expression compared with control by line graph. White dotted line outlining the frontal bone primordia by morphology. Scale bar=100μM.
